## Supplemental Table S2 for "Exercise-induced changes in climbing performance"

**Table S2. Quantitative genetics parameters.**

| Parameter | Symbol | Climbing - control | Climbing - treated |
| --- | --- | --- | --- |
| Mean | μ | 1.522 | 1.524 |
| Genetic variance | σ_G_^2^ | 0.049 | 0.052 |
| Genetic standard deviation | σ_G_ | 0.221 | 0.228 |
| Environmental variance | σ_E_^2^ | 0.031 | 0.030 |
| Environmental standard deviation | σ_E_ | 0.177 | 0.172 |
| Phenotypic variance | σ_P_^2^ | 0.080 | 0.082 |
| Phenotypic standard deviation | σ_P_ | 0.283 | 0.286 |
| Heritability | H_2_ | 0.610 | 0.638 |
| Coefficient of genetic variation | CV_G_ | 14.541 | 14.983 |
| Coefficient of environmental variation | CV_E_ | 11.624 | 11.277 |
| Cross-sex genetic correlation | r_MF_ | 0.834 | 0.763 |
| Genetic correlation | r_g_ | 0.850 | 0.766 |
